# Sex-differentiated hormonal microenvironments recapitulate in vivo liver metabolism in human iPSC-derived organoids

**DOI:** 10.64898/2026.05.09.723948

**Authors:** Stefan Giselbrecht, Rhiannon Grant

## Abstract

Bioengineers strive to recreate *in vivo* microenvironments *in vitro* to reduce our use of animal models and provide insights into human biology. While liver models show promise, sex differences in liver biology remain largely neglected in preclinical studies. Despite the 2014 EU mandate for the inclusion of women in clinical trials, decoupling of research data by sex is historically rare, with only 11% of papers disaggregating data by sex. This gap contributes to women being more susceptible to drug-induced liver injury (DILI) and being underserved in drug development, as well as to costly drug attrition levels. Here we present a novel approach to modelling sex differences in vitro. Human induced pluripotent stem cells (iPSCs) from both male (XY) and female (XX) donors, were differentiated into hepatocyte liver spheroids and exposed to *in vivo*-mimicking levels of testosterone, progesterone, and oestrogen in high-throughput microwell format. We successfully recapitulated sex-specific metabolic profiles and demonstrated significant differences in CYP1A2 and CYP3A4 drug metabolism and gene expression patterns consistent with reported *in vivo* observations, without compromising cell viability. These findings validate the utility of sex-differentiated microenvironments in early-stage research, offering a pathway to refine animal and clinical trials and improve therapeutic outcomes for all sexes.

## 1. Introduction

Translational medicine has been constrained by disconnection between preclinical models and human physiological reality. Animal models and simplified two-dimensional cell cultures frequently fail to recapitulate the complex, dynamic microenvironment of the human body(1–4). Additionally, biomedical research has historically prioritised reduction of variability to achieve statistical power. Male subjects were preferred because they were perceived to eliminate the ‘confounding’ variable of the oestrous cycle(5). This approach introduced systemic bias rendering a vast portion of the human population invisible in foundational drug development data. As few as 11% of biomaterial studies explicitly report the sex of cells or animals used(6), and even fewer disaggregate data to analyse sex-specific effects(7). This limitation is problematic across multiple fields but is particularly pronounced regarding biological sex influences on pharmacokinetics, pharmacodynamics, and disease progression. While the proportion of clinical trial reports incorporating sex-based analysis has risen from 12% to 33% in recent years(8), at least one pharmacokinetic parameter shows a sex difference of 50% or more for many non-oncology drugs(9) and a systematic review identified statistically significant pharmacokinetic differences for 15 commonly used anticancer drugs(10).

The liver serves as the primary site of drug metabolism and detoxification and thus understanding it’s pharmacokinetic behaviour and disease progression is key to understanding drug development and human health. Hepatic function is modulated by sex hormones, producing distinct metabolic phenotypes between males and females(11–13). These differences manifest prominently in cytochrome P450 enzyme expression and activity. CYP3A4, metabolizing approximately 50% of clinically used drugs, is consistently expressed at higher levels in females. Conversely, CYP1A2 typically exhibits higher activity in males(14). These enzymatic variances translate directly into clinical outcomes including adverse drug reactions and drug-induced liver injury, to which women are twice as likely to suffer(15).

Current in vitro liver models do not recapitulate these sex differences in vitro. Two-dimensional liver cell cultures suffer from rapid dedifferentiation and loss of metabolic function(4). Three-dimensional organoids and spheroids have emerged as superior alternatives, but they both often rely on standard culture media lacking hormonal complexity(16–18). Standard media formulations typically contain undefined foetal bovine serum, human albumin or phenol red, introducing uncontrolled oestrogenic activity, or are completely devoid of sex steroids(19). This creates a ‘hormonal desert’ environment failing to drive sex-specific gene expression. In addition, we can derive sex specific organoids via induced pluripotent stem cell (iPSC) technology – providing a renewable sources of patient-specific hepatocytes with XX or XY phenotypes. However, iPSC-derived hepatocytes can retain foetal-like characteristics including low expression of adult CYP enzymes(20). Addition of specific hormonal cues during maturation could serve as powerful drivers of adult-like phenotypes, furthering the model technology and improving our knowledge of CYP metabolism.

This study posits that neglect of sex differences in preclinical liver models is a solvable bioengineering problem. Established, robust 3D cell culture methods(21) enable creation of microenvironments sustaining distinct hormonal conditions allowing simultaneous culture of male and female iPSC-derived liver organoids under controlled conditions mimicking in vivo endocrine landscapes. In this study, we exposed iPSC-derived XX and XY liver organoids to sex-differentiated hormonal microenvironments (containing specific levels of oestradiol, dihydrotestosterone, and progesterone) for seven days and assayed for cell viability, gene expression (CYP1A2, CYP3A4, albumin, UGTB17), and enzymatic drug metabolism activity. Results indicate that these hormonal conditions successfully recapitulated in vivo sex-specific metabolic profiles, with male-derived organoids exhibiting significantly higher CYP1A2 activity and female-derived organoids showing elevated CYP3A4 activity. Furthermore, cross-control experiments demonstrated that these metabolic differences were primarily driven by the hormonal environment rather than genetic background alone, confirming that sex-differentiated microenvironments can effectively model sex-specific drug metabolism without compromising cell viability. By establishing a scalable, simple platform that integrates sex as a biological variable at the earliest preclinical stages, this work offers a direct solution to the historical neglect of sex differences in biomedical research. This approach has the potential to significantly reduce the reliance on animal models, lower costly late-stage drug attrition rates caused by unforeseen sex-specific toxicities, and ultimately mitigate the disproportionate risk of drug-induced liver injury (DILI) in women by enabling earlier identification of sex-dependent pharmacokinetic risks during drug development.

## 2. Results

### 2.1 Recapitulation of sex-specific CYP activity

Exposure to sex-differentiated hormone levels resulted in distinct metabolic profiles (Figure 1). CYP drug-metabolising enzyme induction followed patterns consistent with in vivo expression. CYP1A2 activity was significantly higher in male-derived organoids, while CYP3A4 activity was elevated in female-derived organoids. Cross-control experiments, where female donors were placed in male media and vice versa, confirmed that these differences were driven by hormonal environment rather than genetic background alone.

**Figure 1.**
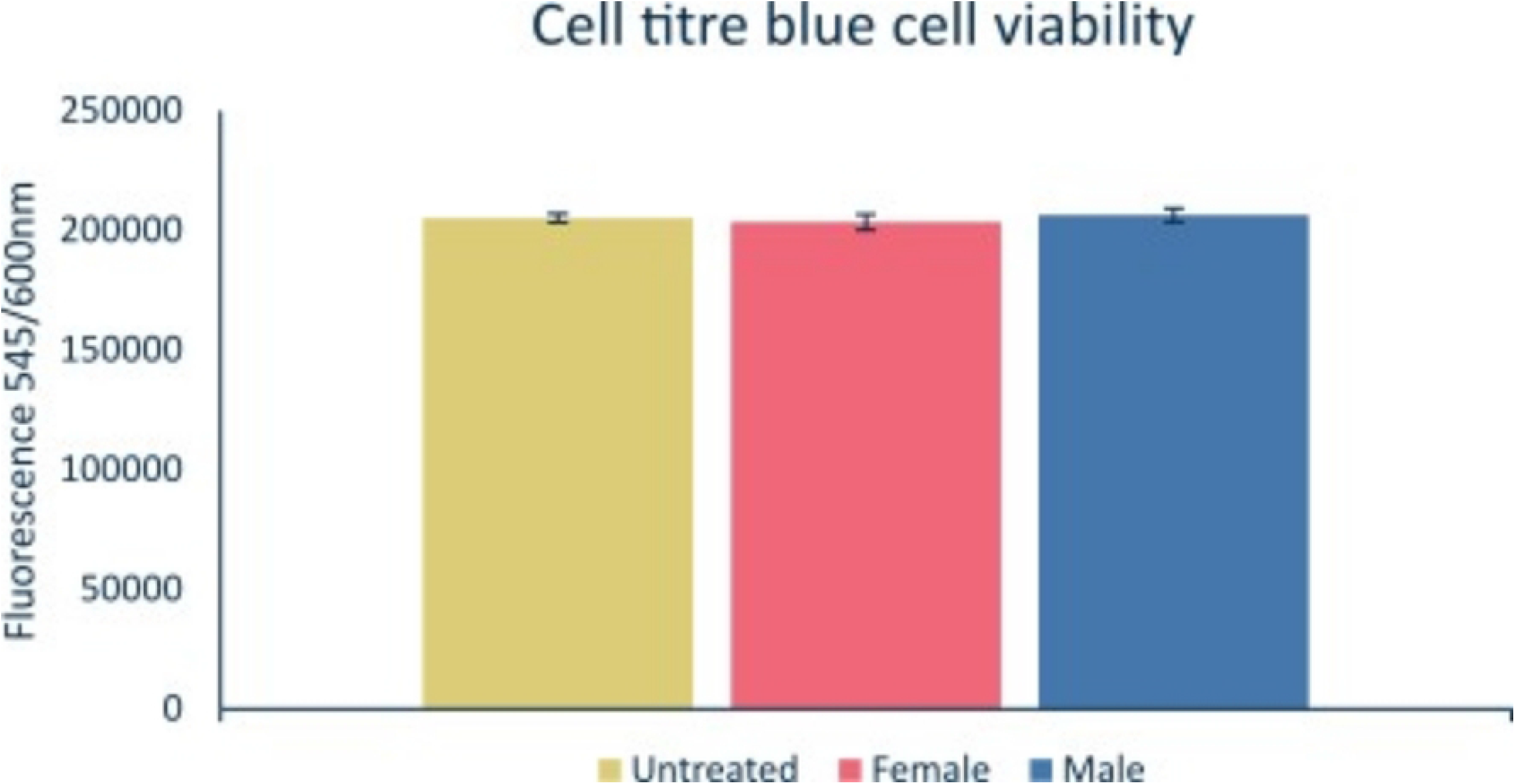
CYP activities following treatments with acetaminophen and erythromycin to induce expression of CYP1A2 and CYP3A4 respectively. N = 6, error bars = SD, one way ANOVA with Games Howell post hoc testing.

**Figure 2.**
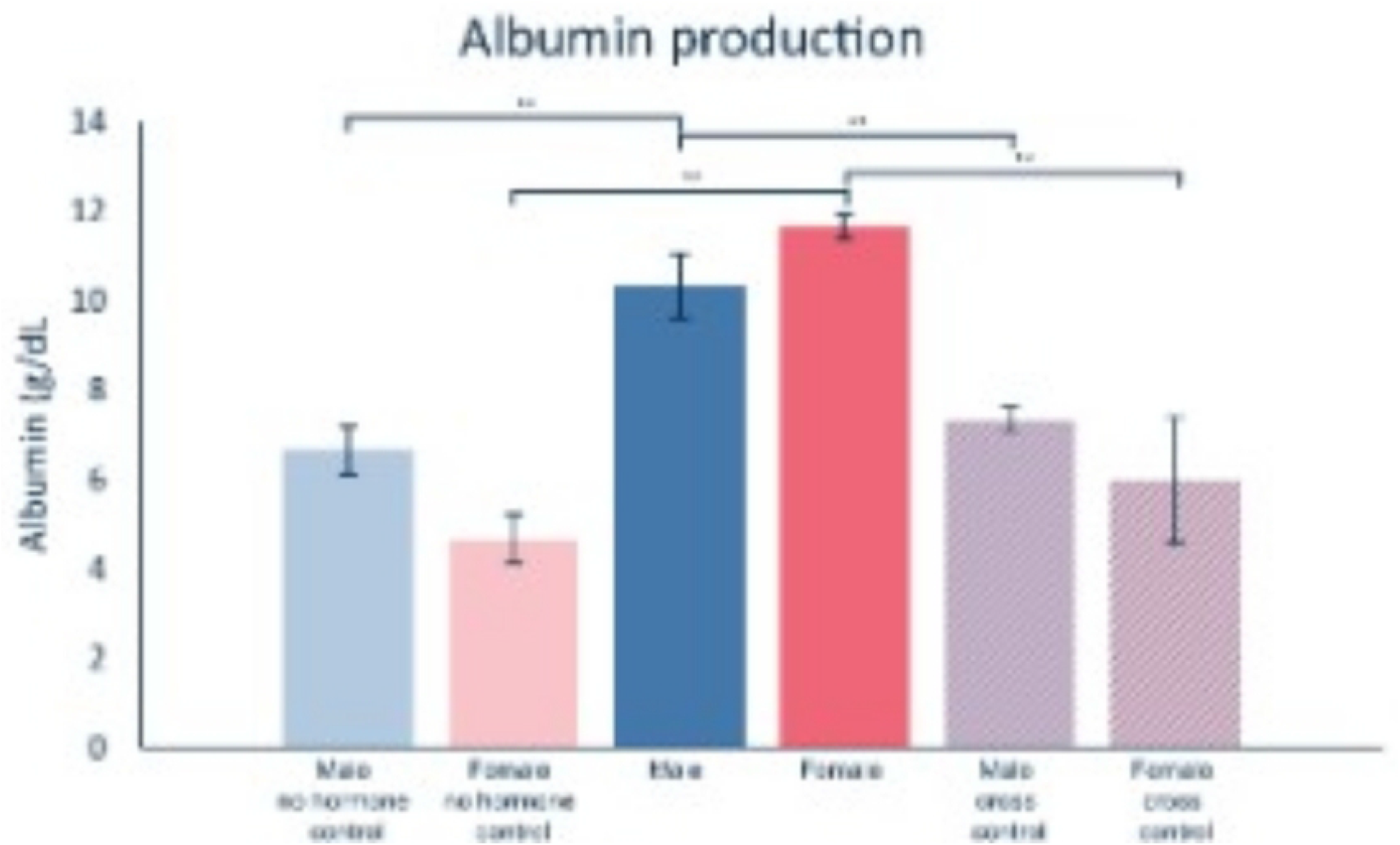
RT-QPCR of key liver function genes CYP1A2 and CYP3A4 (drug metabolising enzymes), Albumin (liver function and protein production), UGTB17 (androgen metabolising enzyme). Fold change, relative to untreated organoids N = 6, error bars = SD, one way ANOVA with Games Howell post hoc testing.

**Figure 3.**
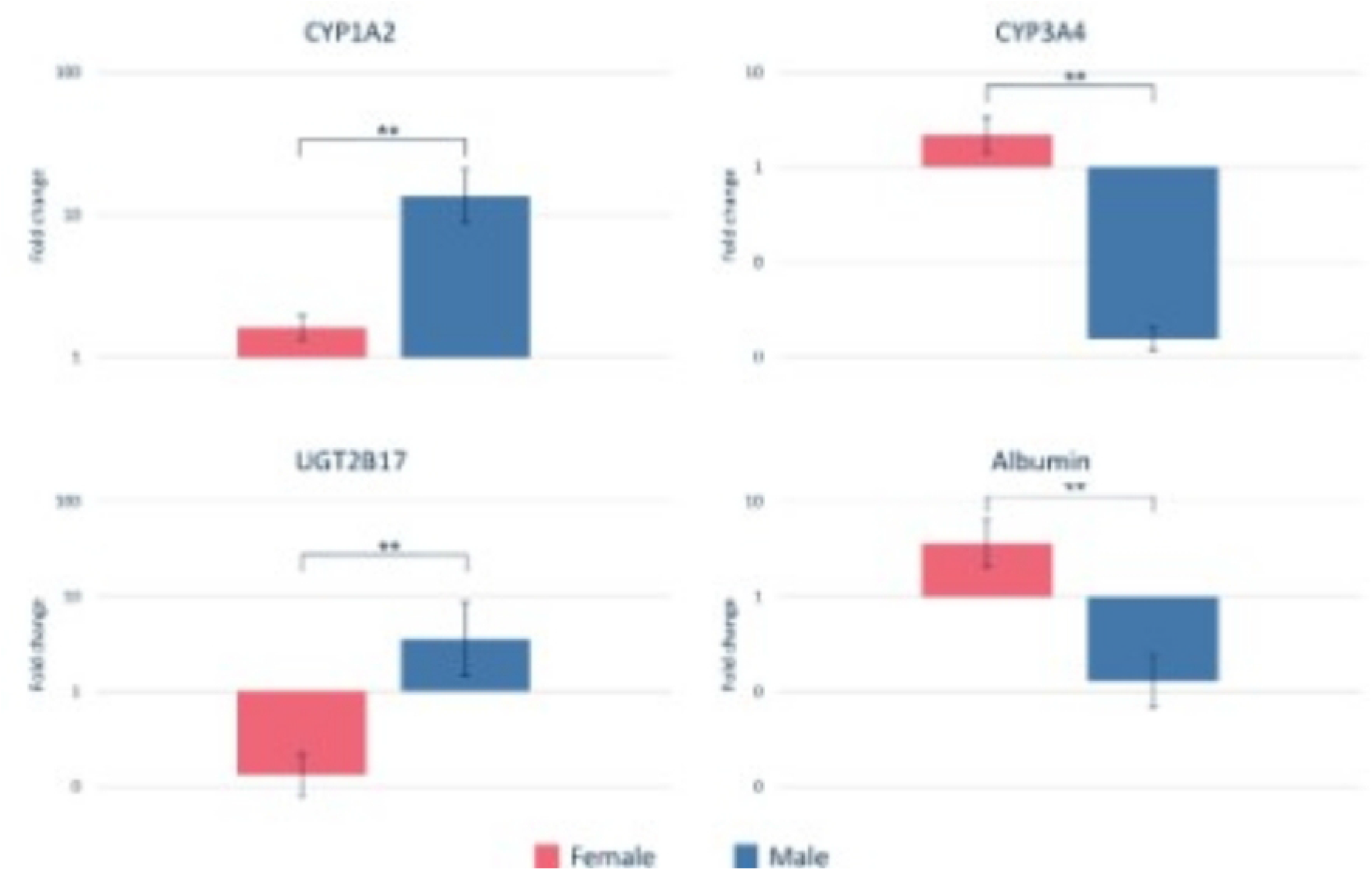
Albumin protein production analysed by bromocresol green assay, as a marker of normal hepatocyte function. N = 6, error bars = SD, one way ANOVA with Games Howell post hoc testing.

**Figure 4.**
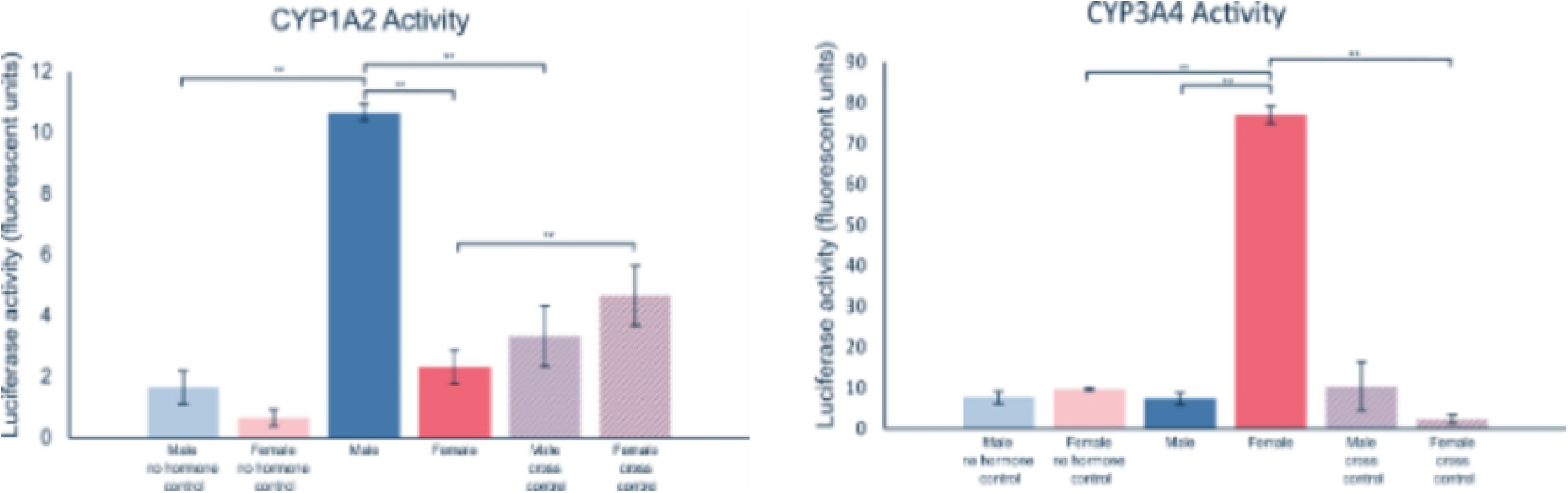
Cell titre blue viability testing of hormonally treated iPSC derived spheroids (female/male) vs untreated spheroids. N = 6, error bars = SD, one way ANOVA with Games Howell post hoc testing.

### 2.2 Gene expression profiles

RT-qPCR analysis revealed significant differences in expression of key liver function genes between male and female organoids. CYP1A2 and CYP3A4 expression levels mirrored enzymatic activity data, showing sex-specific upregulation. UGTB17, an androgen-metabolising enzyme, showed differential expression consistent with hormonal milieu. Albumin gene expression remained robust across all groups, though expression was lower in male organoids, indicating maintained hepatocyte identity.

### 2.3 Functional viability and protein production

Albumin protein production was assessed via bromocresol green assay. Albumin production confirmed that hepatocytes remained functional across all conditions. No significant sex-differences are observed in albumin secretion levels in vivo in humans, serving as a stable baseline for cell health. Albumin production is upregulated significantly by hormonal treatments, demonstrating that sex-differentiated microenvironment supports healthy organoid growth.

### 2.4 Functional viability and protein production

Cell viability was assessed via cell titre blue analysis (Promega) and indicated that the survival of spheroids was not affected by hormonal exposure, bo significant differences were observed between cells treated with no hormones vs those treated with female vs male hormones indicating that other results are not caused by reduced viability.

## 3. Discussion

This work demonstrates succinctly that sex-differentiated hormonal microenvironments can effectively recapitulate in vivo sex-specific metabolic phenotypes in human iPSC-derived liver organoids. Our findings show that exposing male (XY) and female (XX) organoids to physiologically relevant concentrations of oestradiol, dihydrotestosterone, and progesterone for seven days is sufficient to drive distinct expression and activity profiles of CYP1A2 and CYP3A4. Specifically, male-derived organoids exhibited significantly higher CYP1A2 activity, while female-derived organoids showed elevated CYP3A4 activity. These results align with established in vivo literature, where CYP1A2 is typically more active in males and CYP3A4 in females, enzymes responsible for the metabolism of approximately 15% and 50% of clinically used drugs, respectively(22,23). Crucially, our cross-control experiments, wherein female donors were cultured in male hormonal media and vice versa, confirmed that these metabolic differences are driven primarily by the hormonal environment rather than immutable genetic background alone. This finding underscores the plasticity of hepatic metabolic phenotypes and validates the hypothesis that the ‘hormonal desert’ of standard culture media is a driver of the loss of sex-specificity in current in vitro models.

The implications of these findings are profound for the field of translational medicine and drug safety. Historically, the neglect of sex as a biological variable in preclinical research has contributed to a significant gap in understanding sex-specific pharmacokinetics and pharmacodynamics. Few studies explicitly report the sex of cells or animals used, and even fewer sex-disaggregate data and results to provide insight into sex differences(6,8,13,15). This omission has real-world consequences: women are twice as likely to suffer from drug-induced liver injury (DILI) and often experience adverse drug reactions due to unaccounted-for metabolic differences(24). By integrating sex-differentiated hormonal cues into early-stage organoid culture, this study offers a scalable solution to identify sex-dependent toxicities and efficacy issues before costly clinical trials. The ability to detect these differences in a high-throughput format could significantly reduce the reliance on animal models, which often fail to predict human sex-specific responses due to species differences in hormone metabolism and receptor expression(25,26).

Furthermore, our data challenges the traditional view that iPSC-derived hepatocytes are inherently limited to a foetal-like state with low CYP expression. While it is well-documented that standard differentiation protocols often result in immature hepatocytes with reduced metabolic capacity(20) our results indicate that the addition of specific hormonal cues can assist in driving more adult-like phenotypes. The upregulation of CYP enzymes in response to sex hormones suggests that the endocrine environment is a critical, yet often overlooked, maturation factor in hepatocyte differentiation. This aligns with studies suggesting that hormonal signalling is essential for the full functional maturation of liver organoids(27). The observation that albumin production remained robust and was even upregulated by hormonal treatments further confirms that the sex-differentiated environment supports, rather than compromises, general hepatocyte health and function.

However, several limitations must be acknowledged. The most significant constraint of this study is the use of a limited number of donor lines (one male and one female). While this approach was necessary to rapidly communicate the proof-of-concept that hormonal microenvironments can drive sex-specificity, it limits the generalizability of the findings to the broader population when knowledge exists that genetic variability among donors can influence basal CYP expression and response to hormonal stimuli(22). Nevertheless, the consistency of the results across the cross-control experiments provides strong evidence that the observed effects are driven by the experimental manipulation (hormones) rather than donor-specific idiosyncrasies. Future studies will be required to incorporate a larger cohort of donors to establish the robustness of this platform across diverse genetic backgrounds, as well as more complex hormonal environments such as the oestrous cycle or circadian signalling.

Additionally, while the seven-day exposure period was sufficient to induce significant changes, the long-term stability of these sex-specific phenotypes remains to be determined. It is unknown whether these metabolic profiles are maintained over extended culture periods or if they revert in the absence of continuous hormonal stimulation. The use of a static hormone concentration also simplifies the complex pulsatile nature of in vivo hormone secretion, particularly for progesterone and oestradiol, which fluctuate during the menstrual cycle(28). Future iterations of this model could incorporate dynamic hormone delivery systems to better mimic physiological rhythms.

Despite these limitations, the current data provides a compelling argument for the adoption of sex-differentiated culture conditions in preclinical liver modelling. The simplicity of the approach-adding defined concentrations of hormones to existing media-makes it highly accessible for widespread adoption in academic and industrial settings. By addressing the historical bias in preclinical models, this work paves the way for more inclusive and predictive drug development pipelines, ultimately aiming to improve therapeutic outcomes for all sexes.

## 4. Materials and Methods

### 4.1 Cell sources and maintenance culture

Human induced pluripotent stem cells derived from male (XY) and female (XX) donor sources were obtained from Leiden University Medical Centre. Lines LUMCi029-A (XY) and LUMC0099iCTRL04 (XX) were maintained on vitronectin in TESR-PLUS media using standard protocols for 7 days prior to spheroid generation and differentiation.

### 4.2 Spheroid generation and hepatocyte differentiation

iPSCs were differentiated into hepatocyte-like cells using established protocols to generate liver organoids(29–33). Differentiation was conducted in non-adherent 96 well round bottom culture plates coated with 5% w/v F-108 pluronic for 1hr at 37°C. Briefly, cells were enzymatically detached from their maintenance culture plates using TrypleE and reseeded into coated 96 well plates at 3000 cells per well before being taken through differentiation protocols as previously published.

### 4.3 Hormone exposure

Liver organoids were cultured in three distinct media conditions. Control media consisted of: Hepatocyte maintenance media containing no phenol red, no serum, and ethanol as hormone diluent control. Male media was hepatocyte media supplemented with 20 pg/ml oestradiol (E2), 0.6 ng/ml dihydrotestosterone (DHT), and 0.5 ng/ml progesterone, and female media was hepatocyte media supplemented with 250 pg/ml E2, 0.1 ng/ml DHT, and 10 ng/ml progesterone. Organoids were exposed to these hormone levels for seven days prior to analysis.

### 4.4 CYP1A2 and CYP3A4 enzymatic activity

CYP1A2 and CYP3A4 enzymatic activities were quantified using the CYP1A2-Glo™ and CYP3A4-Glo™ Assay systems (Promega Corporation, Madison, WI, USA), which utilise luminescent reporter gene constructs to measure enzyme induction. Following the seven-day hormonal exposure period, liver organoids were washed three times with phosphate-buffered saline (PBS) and lysed directly in the culture wells according to the manufacturer’s instructions.

Acetaminophen served as the specific substrate and inducer for CYP1A2-mediated metabolism. Organoids were treated with 2 mM acetaminophen (paracetamol) dissolved in the respective sex-differentiated media for 24 hours prior to lysis. Erythromycin was utilized as the selective inducer for CYP3A4. Organoids were exposed to 10 µM erythromycin in the culture media for 24 hours. Control groups were maintained in media containing the vehicle (ethanol) without inducers.

Following the 24-hour induction period, 50 µL of the Luciferase Detection Reagent was added to 50 µL of the cell lysate in a white 96-well plate. The mixture was incubated at room temperature for 10 minutes to allow for the conversion of the luciferin substrate by the luciferase enzyme, which is expressed under the control of the CYP-responsive promoter elements. Luminescence was measured using a plate reader (Clariostar) with an integration time of 1 second per well.

Relative luminescence units (RLU) were recorded and normalized to total protein content, which was determined using the Pierce™ BCA Protein Assay Kit (Thermo Fisher Scientific) on parallel lysates, to account for variations in cell number and viability. Data were expressed as RLU for each sex-specific condition. All assays were performed in triplicate (N=6 biological replicates), and statistical significance was determined using one-way ANOVA with Games-Howell post-hoc testing.

### 4.5 Gene Expression Analysis

Total RNA was extracted from the liver organoids using TRIzol™ Reagent (Invitrogen, Carlsbad, CA, USA) according to the manufacturer’s protocol. RNA concentration and purity were assessed spectrophotometrically (NanoDrop™, Thermo Fisher Scientific). Quantitative PCR was performed using SYBR™ Green Master Mix (Applied Biosystems) on a StepOnePlus™ Real-Time PCR System (Applied Biosystems).

Gene expression levels were quantified for *CYP1A2, CYP3A4, ALB* (albumin), and *UGT2B17*. The housekeeping gene *GAPDH* was used as an internal reference for normalization. Primer sequences were designed to span exon-exon junctions to avoid genomic DNA amplification (sequences available upon request). Relative gene expression was calculated using the 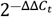method. All reactions were performed in technical triplicates, and data were analyzed from six independent biological replicates (N=6). Statistical differences in gene expression between sex-specific and hormonal conditions were evaluated using one-way ANOVA with Games-Howell post-hoc testing.

### 4.6 Albumin production

Albumin secretion levels were quantified to assess hepatocyte functional maturity and viability using the Bromocresol Green (BCG) assay. Following the seven-day hormonal exposure and induction periods, conditioned media was collected from the organoid cultures and centrifuged at 300 × g for 5 minutes to remove cellular debris.

For albumin measurement, 10 µL of the clarified conditioned media was mixed with 200 µL of BCG working solution according to the manufacturer’s instructions. The mixture was incubated at room temperature for 5 minutes to allow the formation of the albumin-BCG complex. Absorbance was measured at 630 nm using a microplate reader (Clariostar).

A standard curve was generated using serial dilutions of human serum albumin (HSA) ranging from 0 to 200 µg/mL in the respective culture media. Albumin concentrations in the samples were interpolated from the standard curve. All assays were performed in triplicate across six independent biological replicates (N=6). Data were expressed as mean ± standard deviation (SD), and statistical comparisons were conducted using one-way ANOVA with Games-Howell post-hoc testing.

### 4.7 Cell viability

Cell viability was assessed to ensure that the sex-differentiated hormonal microenvironments did not induce cytotoxicity during the seven-day culture period. Following the final media exchange, 20 µL of CellTiter-Blue® Reagent (Promega) was added directly to 100 µL of culture media, resulting in a final reagent concentration of 10% (v/v). The plates were incubated at 37°C in a humidified atmosphere containing 5% CO_2_ for 3 hours to allow for the reduction of resazurin to the fluorescent resorufin by metabolically active cells.

Fluorescence intensity was measured using amicroplate reader (Clariostar, excitation wavelength: 560 nm; emission wavelength: 590 nm). Background fluorescence from media-only controls (without cells) was subtracted from all sample readings. All assays were performed in technical triplicates across six independent biological replicates (N=6). Data were presented as mean ± standard deviation (SD), and statistical significance between groups was determined using one-way ANOVA with Games-Howell post-hoc testing.

## 5. Conclusion

In conclusion, this study demonstrates that sex-differentiated hormonal microenvironments can successfully recapitulate in vivo sex-specific metabolic profiles in human iPSC-derived liver organoids. By exposing male and female organoids to physiologically relevant levels of oestradiol, dihydrotestosterone, and progesterone, we observed a significant divergence in CYP1A2 and CYP3A4 activity and expression, mirroring established clinical observations. The cross-control experiments confirmed that these differences are driven by the hormonal environment, highlighting the plasticity of hepatic metabolism and the critical role of endocrine cues in maintaining sex-specific phenotypes in vitro.

While the current study utilises a limited number of donor lines to rapidly establish proof-of-concept, the results provide a robust foundation for the integration of sex as a biological variable in early-stage drug development. This approach offers a practical, scalable, and cost-effective solution to the historical neglect of sex differences in preclinical research. By enabling the early identification of sex-dependent pharmacokinetic risks and toxicities, this model has the potential to reduce drug attrition rates, minimize the reliance on animal models, and ultimately mitigate the disproportionate risk of adverse drug reactions in women. Future work will focus on expanding the donor pool to assess genetic variability, investigating the long-term stability of sex-specific phenotypes, and exploring the application of this platform to other tissue types and disease models. The adoption of such sex-inclusive strategies is essential for advancing precision medicine and ensuring that therapeutic interventions are safe and effective for the entire population.

## 6. Acknowledgements

This work was performed at the MERLN Institute for Technology-Inspired Regenerative Medicine at Maastricht University and the Institute for Bioengineering (IBioE) at the School of Engineering, University of Edinburgh. The authors gratefully acknowledge the technical assistance provided by Barry Jutten and Whitney Van der Vin, whose expertise was invaluable to the execution of these experiments. This research was supported by the University of Edinburgh Chancellor’s Fellowship awarded to R.G. and the NWO Open Mind Grant (No. 19553), also awarded to R.G.

## 7. Author contributions

R.G. conceived the project, secured funding, performed the experimental work, and drafted the manuscript. S.G. provided supervision, and contributed to the conceptual design and revision of the manuscript. Both authors reviewed and approved the final version of the paper.

## 8. Data availability statement

The data that support the findings of this study are available from the corresponding author upon reasonable request.

## 9. Additional information (competing interests)

The authors declare no competing interests.

